# Multiplexed protein detection and parallel binding kinetics analysis with label-free digital single-molecule counting

**DOI:** 10.1101/2022.06.08.495341

**Authors:** Xinyu Zhou, Rui Wang, Zijian Wan, Pengfei Zhang, Shaopeng Wang

## Abstract

Multiplexed protein detection is critical for improving the drug and biomarkers screening efficiency. Here we show that multiplexed protein detection and parallel protein interaction analysis can be realized by evanescent scattering microscopy with label-free digital single-molecule counting. We implemented an automatic single-molecule counting strategy with high temporal resolution to precisely determine the binding time, which improves the counting efficiency and accuracy. We show that digital single-molecule counting can recognize proteins with different molecular weights, thus making it possible to monitor the protein binding processes in the solution by real time tracking the numbers of free and bound proteins landing on the sensor surface. Furthermore, we show that this strategy can simultaneously analyze the kinetics of two different protein interaction processes on the surface and in the solution. This work may pave a way to investigate complicated protein interactions, such as the competition of biomarker-antibody binding in biofluid with biomarker-protein binding on the cellular membrane.

## Introduction

Protein plays a key role in the structures and activities of living systems. Determining the proteins and their interactions is critical for drug screening^1,2^, clinical biomarker analysis^3,4^, and understanding the biological processes at the molecular level^5,6^. Diverse technologies have been developed for protein analysis, including mass spectrometry^7^, enzyme-linked immunosorbent assay^8^, western blot^9^, and surface plasmon resonance (SPR)^10,11^. However, these traditional techniques usually only provide ensemble measurement results and lack the capability to analyze the highly heterogeneous proteins at a level of detail. In the recent decade, label-free single-molecule imaging approaches have been developed to push beyond ensemble averages and conduct the statistical analysis of intrinsic protein properties such as molecular weight and binding processes. These techniques include interferometric scattering microscopy (iSCAT)^12–14^, photothermal microscopy^15,16^, and recently developed plasmonic scattering microscopy (PSM)^17– 20^ and evanescent scattering microscopy (ESM)^21^.

To precisely analyze the single-molecule signal, which is usually very weak, these imaging-based approaches usually need to remove the strong but static backgrounds with differential processing, namely subtracting or dividing one frame by the previous frame. Meanwhile, the raw image frames are usually required to be averaged at the temporal domain to suppress the shot noise. Unfortunately, the protein binding events are random in the time domain. The extracted single molecule signal intensity depends on the relative time position of the binding events in the averaged frames. The molecules bind to the sensor surface at around the middle of one average period, will have underestimated image intensity or missed if the image processing does not consider the time point of the binding events. This measurement error will limit the system precision in recognizing the proteins with different molecular weights in a mixed sample and in analyzing the kinetics of binding processes, which relies on digitally counting the binding events over time.

Here, we present an automatic single-molecule counting algorithm for ESM that uses moving average and time tracking approaches to determine the time point of binding events with the temporal resolution of the camera exposure time. It has been shown that the moving average is important for obtaining the single-molecule signals precisely in iSCAT^12,22^. By processing the same experimental data, this algorithm shows that it misses few events and provides more precise image intensity than the classical algorithm. Next, we show that this algorithm can recognize proteins with different molecular weights, thus allowing for monitoring the kinetics of protein binding in solution phase by tracking the numbers of free protein and bound protein complex hitting on the sensor surface, providing one approach for multiplexed protein analysis. Furthermore, we show that this strategy can simultaneously analyze the kinetics of two protein interaction processes on the surface and in the solutions, providing one approach for parallel binding kinetics analysis.

## Results

### Algorithm principle

ESM system is employed in this study. The 450 nm laser is conditioned to illuminate one indium tin oxide (ITO) coated cover glass with an incident angle of ∼65° through a 60x objective. The incident angle is larger than the critical angle to excite evanescent waves on the surface (Figure 1a)^21^. A 50x objective is placed on the top of ITO-coated cover glass to collect the evanescent waves scattered by proteins and surface roughness to form the ESM images.

**Figure 1.**
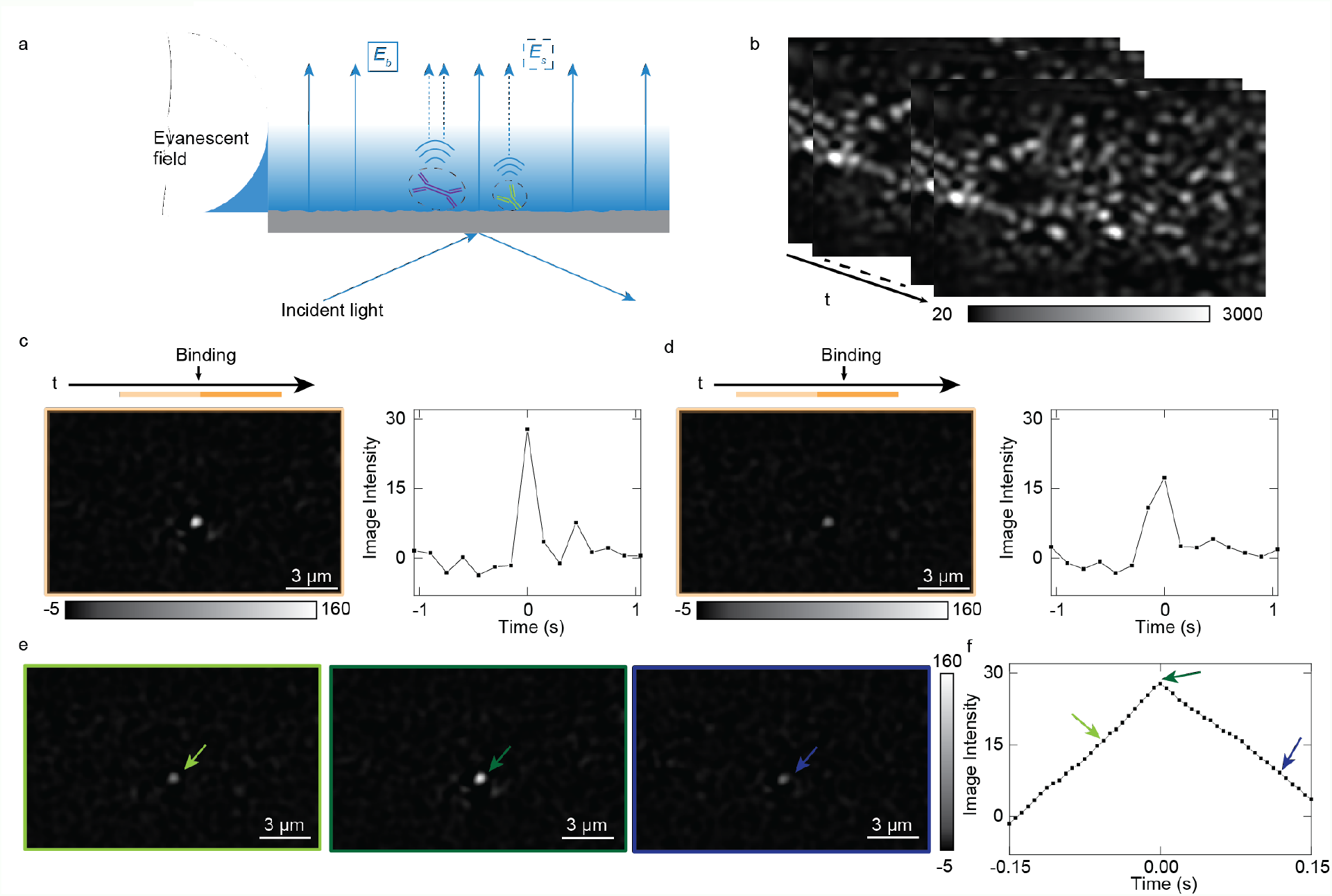
System and image processing principles. (a) The ESM schematic, incident light creates an evanescent field on the surface, and scattering light of protein (E_s_) and glass surface (E_b_) are collected by the top-mounted objective (not shown). (b) ESM raw image sequence. (c) Single protein image after data processing with the correct estimation of the protein binding time in (b), where the light and dark, orange-colored bar above the image indicate the selected time zone for the single protein analysis. The binding time is in the middle of two image stacks. The differential between averaged images from the dark zone and light zone reveals the protein binding. (d) Single protein image after data processing with the wrong estimation of the protein binding time, leading to miscalculation of image intensity. By moving-average the raw image sequence with single frame steps, a protein binding event obtained from the differential of the two averaged images will have an intensity profile that firstly increases and then decreases (e, f), arrows in the (f) correspond to the images in the (e) with the same color.

To suppress the shot noise, the raw images recorded at 160 frames per second (fps) are firstly averaged over time, where the average period is configured to be ∼150 ms (Figure S2). Traditionally, the binding image was obtained by implementing a differential process subtracting the previous frame from the present frame, where a binding event appears as a bright spot (Figure 1c). When a protein bind to the surface, all pixel values within the Airy disk pattern are summed up at the binding site as the image intensity^18^. However, this approach is usually accompanied with image intensity fluctuations over binding time because the protein binding event usually does not ideally happen in the middle of two consecutive averaging stacks, leading to inaccurate estimation of the single-molecule signal intensities (Figure 1d).

To determine the single-molecule signal more accurately, we scan the binding event in the raw image sequence with a 150 ms moving average window at a single frame step (6.25 ms) to precisely estimate the binding time. This frame step is fixed for the following experiments except else mentioned. Starting with the first frame, we average the two sequential image stacks (frames 1 to n and frames n+1 to 2n) without image overlapping to generate two averaged images. The averaging is performed for each pixel in the time domain with an average period of 150 ms. Next, we subtract the first averaged image from the second averaged one, obtaining the differential image. The signal change, such as a binding or unbinding event, will be shown in a differential image as a bright or dark spot, respectively. Finally, we repeat the above steps sequentially on every subsequent frame in the raw image sequence. For a protein binding to the surface event, the moving-averaging-and-differential process will generate an increase-then-decrease intensity profile in the binding site (Figure 1e, f) with temporal resolution equal to the duration of single frame exposure time. The maximum intensity in the middle of the intensity profile is the true protein intensity, which exactly locates the binding time in the middle of two averaging stacks.

### System Calibration

IgG (150kDa), IgA (385kDa), and IgM (950kDa) were measured separately in the experiment to calibrate the ESM system. Protein in phosphate-buffered saline (PBS) buffer was flowed over the chip. The raw image sequence was recorded over time for each measurement (Method). Figure 2a-c shows protein binding images after moving-averaging and differential data processing. We constructed the histograms by tacking each protein binding event and obtained each protein intensity. Gaussian fitting was applied to the histogram to obtain the mean intensity of the protein. We received 867 binding events with ∼21% higher protein intensities and ∼125% more binding numbers than data processing without time-scanning (Figure S3). After getting the mean intensity of each protein type, a calibration curve of protein mass versus image intensity was made. As expected, the calibration curve reveals a linear relationship between protein mass and image intensity.

**Figure 2.**
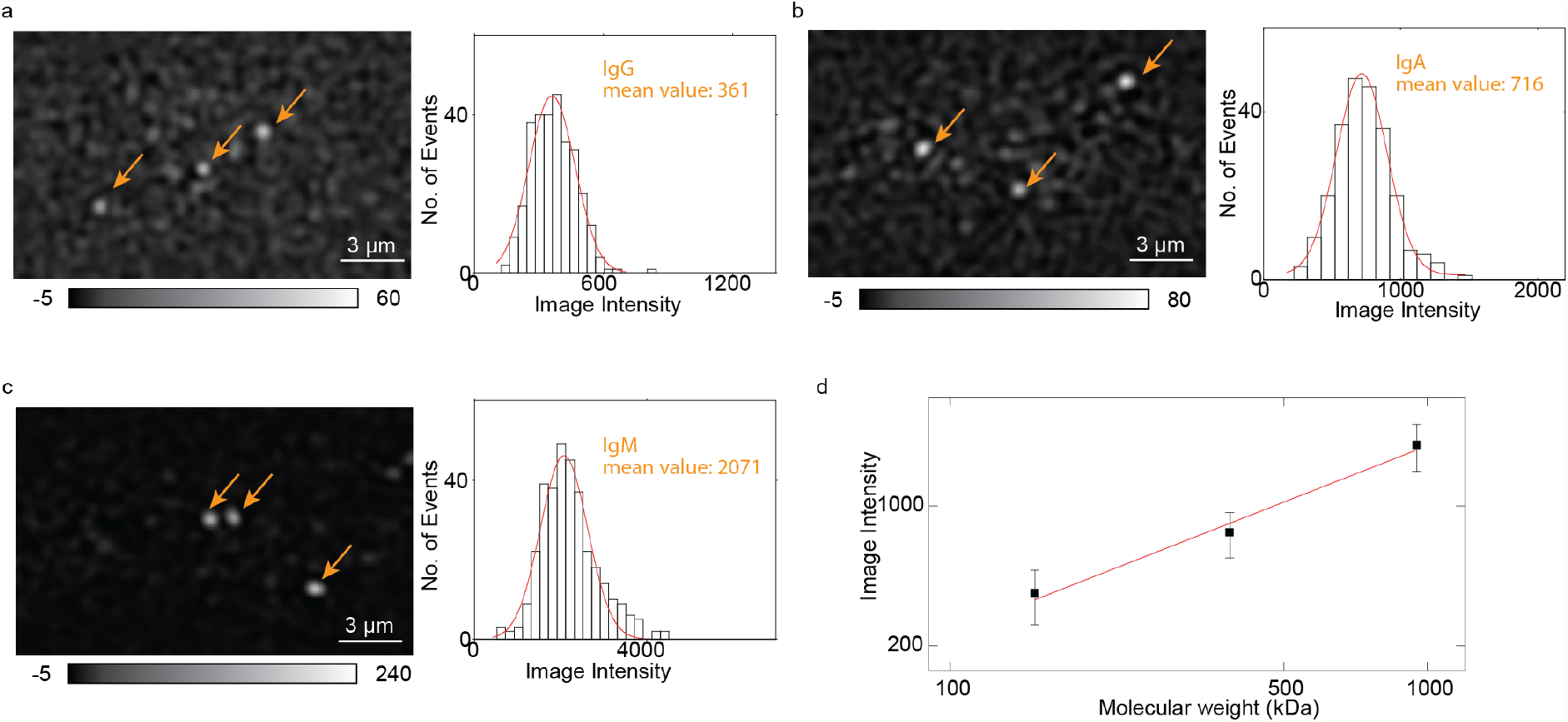
System calibration. (a-c) IgG, IgA, and IgM images (after image processing) and intensity histogram. Orange arrows indicate the binding spots. The red curve is the Gaussian fitting. (d) ESM intensity versus protein mass. The image intensities are obtained from the histograms of each protein in (b-d). The error bars indicate the standard deviation of Gaussian fitting.

### Binding kinetics of IgM to anti-IgM on the surface

To demonstrate the capability of ESM measuring protein binding kinetics with antibodies surface-immobilized approach, we performed IgM and anti-IgM binding kinetics measurement with anti-IgM modified on the surface. We first flowed 2.5 nM IgM solution over the chip for IgM association with the antibodies immobilized on the surface, then flowed PBS buffer over the surface to allow the dissociation of IgM. We tracked the binding and unbinding events of the IgM in real time and plotted the bound IgM numbers versus time to form the binding kinetics curves^17,18^ (Figure 3b). The fitting of the curves with the first-order binding kinetics model determines the association (*k*_on_) and dissociation (*k*_off_) rate constants, which are 1.66 × 10^6^ M^−1^ s^−1^ and 9.46 × 10^−4^ s^−1^, respectively. From *k*_on_ and *k*_off_, the equilibrium dissociation constant (*K*_D_ = *k*_off_ / *k*_on_) is determined to be 570 pM. These values are in good agreement with the results measured with the ensemble SPR (Figure S4). We collected individual protein binding intensity and made the histogram. The mean intensity of binding proteins is consistent with what we obtained in the calibration experiment (Figure 3c), confirming the single IgM detection.

**Figure 3.**
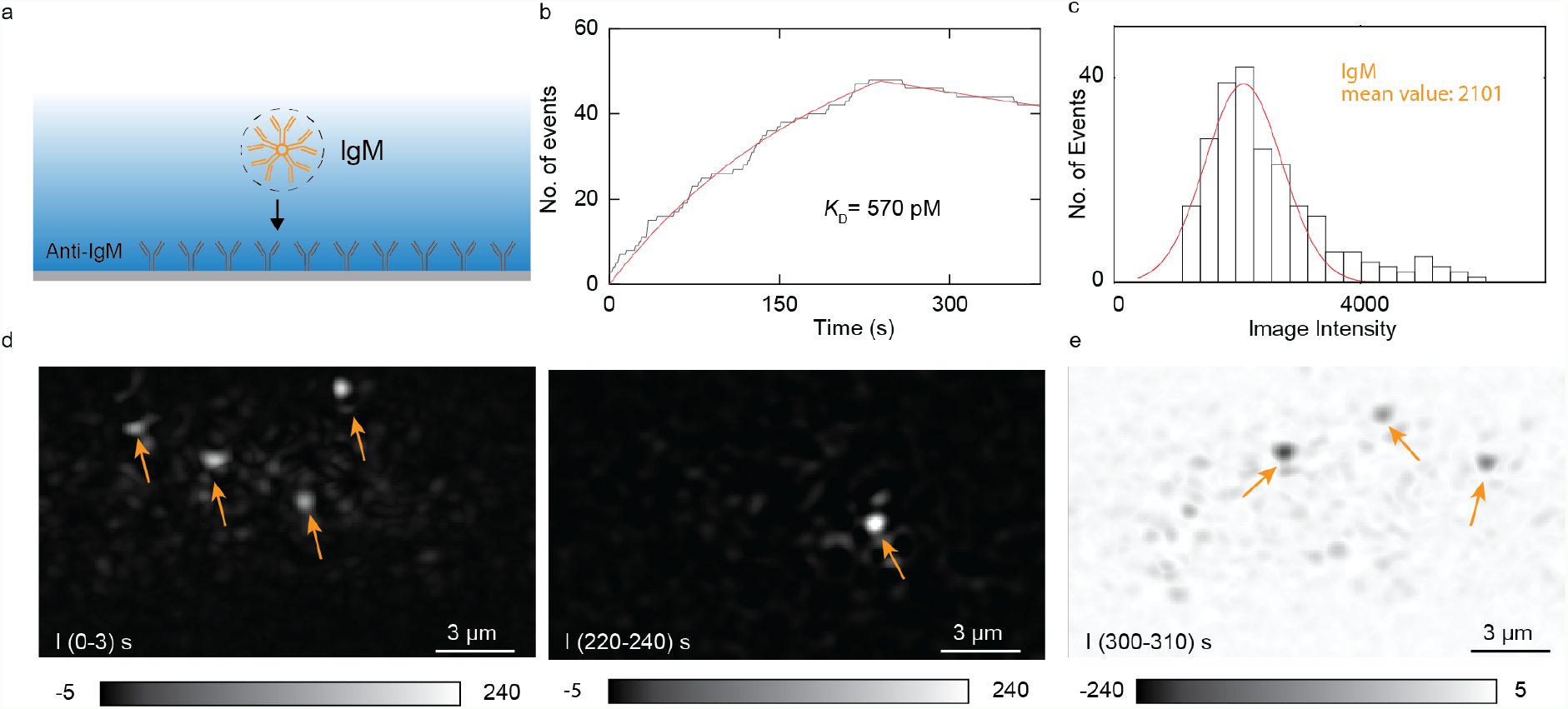
Analysis of IgM binding to anti-IgM on the surface. (a) Schematic of the experiment. (b) Accumulated counts of bound IgM molecules (black line) and first-order binding kinetic fitting (red line). (c) Histogram of IgM binding intensity. Gaussian fitting (red curve) shows the distribution and mean intensity. The histogram is obtained from 3 experiments. (d) Differential images show IgM binds to the surface at two different times. Arrows point to binding IgM spots. (e) IgM unbinds from the surface. Arrows point to unbinding spots.

### Binding kinetics of IgA to anti-IgA in free solution

We demonstrate that ESM can measure free solution protein-protein interaction through counting the protein landing events to the sensor surface. Considering the process of two monomer species forming the complex in the solution, over time, the concentration of monomers will decrease, and the concentration of the complex will increase. By simultaneously counting the random landing events of the monomer and the complex to the surface, we disclosed the free monomer ratio as a function of time in the solution and calculated the binding kinetics. In the experiment, we modified the ITO surface with bovine serum albumin (BSA), which is a commonly used surface blocker. In this way, the probability of protein landing on the surface scales with the concentration of the protein in the solution. The protein solution was prepared with 5 nM anti-IgA mixed with 5 nM IgA. Then the protein solution was flowed over the BSA chip immediately after the mixing. We tracked the protein landing events in real time. Anti-IgA, IgA, and complex are distinguished based on their image intensity, and their counts are accumulated over time separately. The IgA monomer solution abundance was calculated from the protein counting numbers (Method). During the experiment, few complexes were detected in the first minute (Figure 4c), and the monomer ratio was close to one. In the third minute, more complexes were detected, and the monomer ratio was close to 0.5 (Figure 4d). We chose a one-minute time window for accumulating the count number and a five-second moving step. The solution abundance of IgA monomer was calculated as a function of time based on counts. The ratio of monomer IgA versus time is shown in Figure 4e. The solid curve shows the fitting result of the monomer ratio decay (Method). We obtained *K*_D_ = 839 pM from the fitting, which is in good agreement with our previous experiment^17^, with *k*_off_ = 1.73 × 10^−3^ s^-1^ and *k*_on_ = 2.18 × 10^6^ M^-1^s^-1^.

**Figure 4.**
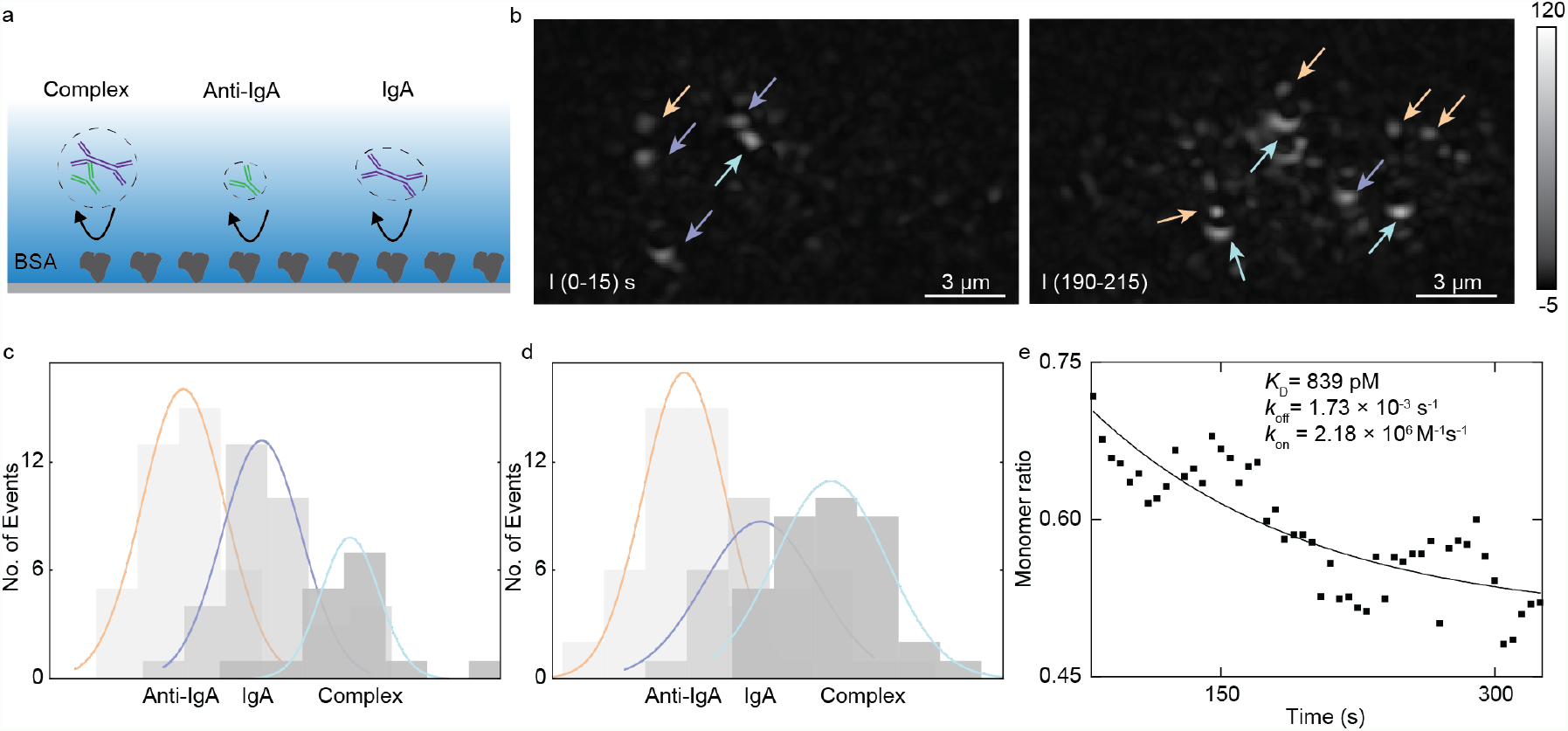
Detection of anti-IgA and IgA interaction in the free solution through surface nonspecific bindings. (a) Schematic of nonspecific binding of anti-IgA, IgA and their binding complex, the surface is modified with BSA. (b) Protein binding image of anti-IgA (orange arrow), IgA (purple arrow), and complex (light green arrow) at different times. Histograms of the first minute (c) and the third minute (d) protein counts accumulated in one minute. Gaussian fitting colors correspond with those in (b). (e) IgA monomer ratio (*X*_IgA_) as the function of time, the solid line is the exponential decay fitting result. Each data point is the ratio in a one-minute time window and separated by five seconds.

### Parallel binding kinetics analysis

The previous sections show that measuring protein-protein interaction is possible either by immobilizing one of the binding antibodies on the surface or by detecting solution protein abundance changes using ESM. Here, we demonstrate that these two methods are compatible in ESM measurement. We simultaneously measured two pairs of protein interactions (anti-IgA to IgA, anti-IgM to IgM) from a protein mixture. Anti-IgA, IgA, and IgM were mixed with 2.5 nM final concentration for each protein in PBS and immediately flowed over the chip. The ITO chip was pre-modified with anti-IgM on the surface. Figure 5b shows IgM bound numbers along the time. IgM dissociates from the surface, providing bound number decay in Figure 5b after PBS was flowed over the chip surface. The fitting of the curves with the first-order binding kinetics model determines the association (*k*_on_) and dissociation (*k*_off_) rate constants, which are 5.21 × 10^6^ M^−1^ s^−1^ and 2.96 × 10^−3^ s^−1^, respectively. From *k*_on_ and *k*_off_, the equilibrium dissociation constant (*K*_D_ = *k*_off_/*k*_on_) is determined to be 568 pM. These values are in good agreement with the stand-alone ESM results shown in Figure 3. At the same time, Anti-IgA, IgA, and their complex hit the surface nonspecifically. We counted protein collision events in real time and calculated the monomer ratio as the function of time (Method). Figure 5c shows the monomer IgA ratio versus time during the experiment. At the start of the protein detection, we had the highest monomer ratio, and in the flowing three minutes this ratio decreased, which is the same as Figure 4. The fitting result of ratio decay is shown in a solid line obtaining *k*_off_ = 1.48 × 10^−3^ s^-1^ and *k*_on_ = 8.81 × 10^5^ M^-1^s^-1^, which leads to *K*_D_ = 1.84 nM.

**Figure 5.**
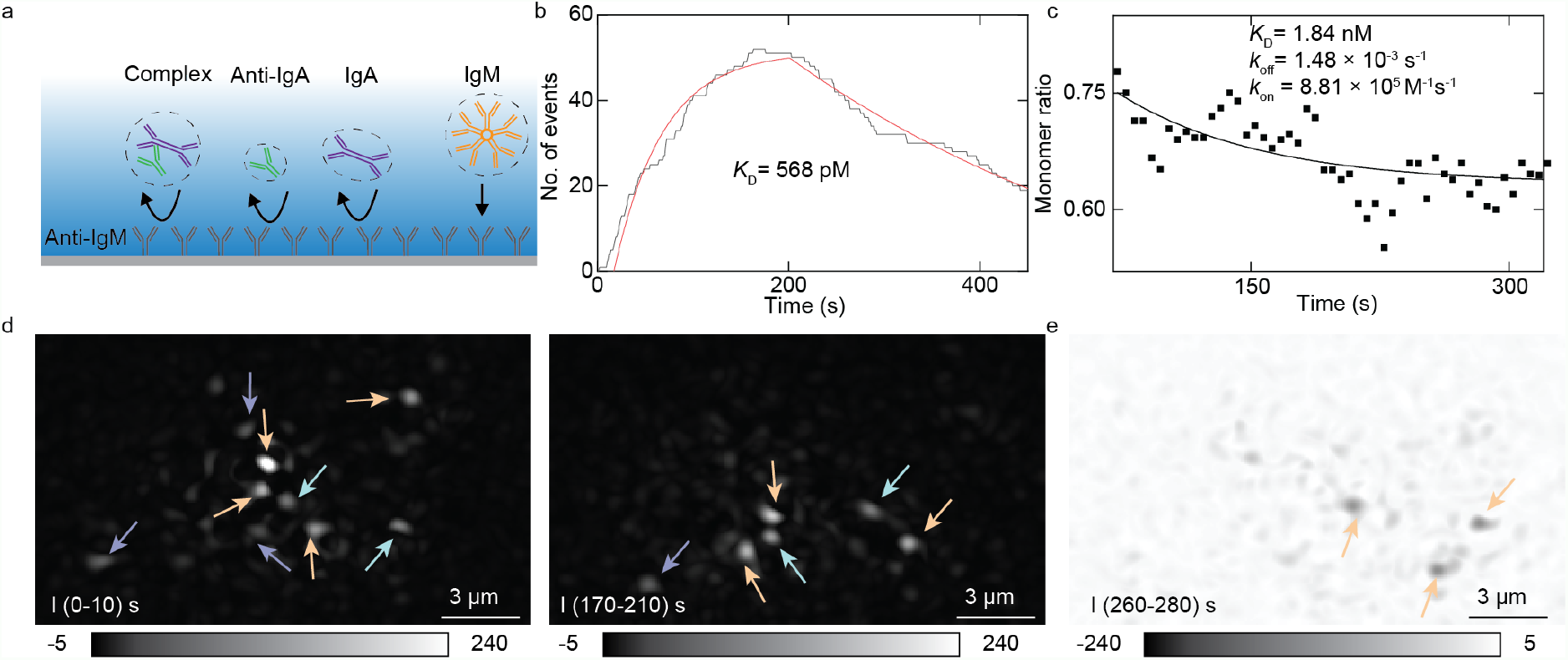
Parallel binding kinetics analysis of anti-IgA, IgA, anti-IgM, and IgM. (a) Schematic of protein binding onto the surface. Anti-IgA, IgA, and their complex bind to the surface nonspecifically. IgM bind to the surface specifically. (b) Count-based anti-IgM and IgM binding interaction measurement, the red curve is the fitting result. (c) The solid line is the monomer decay fitting result of anti-IgA and IgA interaction measurement. (d) IgA (purple arrow), complex (light green arrow), and IgM (orange arrow) bind to the surface at different times. (e) IgM unbinds from the surface. Arrows indicate the unbinding spots.

## Discussion

The single-molecule imaging systems can measure single protein size and number separately, which is a distinct advantage over ensemble molecular interaction analysis. The ensemble approaches only give the integrated results of both quantities, which are not only sensitive to impurity but also impossible to measure multiple protein species at the same time without spatial separation. However, the single-molecule signal is usually weak and challenging to be precisely achieved with conventional approaches. One of the issue is that traditional average-and-differential image processing for shot noise reduction also sacrifices the temporal resolution needed to locate the timing of single molecule binding events. To overcome this issue, we use a moving-average of the images with a single frame step to find the accurate binding time of each protein. In this way, we can count more single protein binding events and determine the single-molecule signals more precisely. Although we use ESM to show the capability of this improved image processing method, the method should be also applicable to PSM and other imaging-based methods that need the average-and-differential image processing approach for shot noise reduction and background subtraction.

To demonstrate the advantage of this improved data processing method, we show that we can track single protein landing events on the surface and distinguish anti-IgA, IgA, and the binding complex based on proteins spot intensities. By counting the ratio of IgA in the total landing events (IgA + complex), we can track the decay of free monomer IgA abundance in the solution during the binding process and obtain the free solution binding kinetics by fitting the decay curve. Furthermore, we can simultaneously count multiple molecules distinct by their intensities to simultaneously analyze the kinetics of binding processes in the solution and on the surface. In the future, we anticipate this capability can help quantitatively estimate the competition between the analyte-protein interaction on the membrane and analyte-antibody binding in the solution.

## Materials and methods

### Materials

ITO chips were purchased from SPI Supplies. Human colostrum IgA was purchased from Athens Research and Technology. Anti-IgA was purchased from BIO-RAD. Phosphate-buffered saline (PBS) was purchased from Corning. PBS was filtered with 0.22 um filters (Millex) before usage. N-ethyl-N’-(dimethylaminopropyl) carbodiimide (EDC) and N-hydroxysulfosuccinimide (Sulfo-NHS) were purchased from Thermo Fisher Scientific. 99.5% Isopropyl Alcohol (IPA) was purchased from Oakwood chemical.

### Experimental Setup

An 80 mW 450 nm laser diode (PL450B, Thorlabs, Newton, NJ, US) is used for creating an evanescent wave in the experiment. The laser diode is mounted to a temperature-controlled mount (LDM38, Thorlabs) and driven by a benchtop diode current controller (LDC205C, Thorlabs) and a temperature controller (TED200C, Thorlabs). Light from the laser diode is conditioned by an achromatic doublet lens group and focused at the back focal plane of a 60x Objective (Olympus APO N 60x Oil TIRF, NA 1.49). The light incident onto the ITO chip with an angle of ∼65° adjusted by a translation stage (XR25P-K2, Thorlabs). The ×50 imaging objective (NA, 0.42) is top-mounted collecting surface, and protein scattered light. A camera (MQ003MG-CM, XIMEA, Münster, Germany) connected to the imaging objective is used to record the scattering images at 160 fps (Figure S1). The image is recorded with a ∼8×10 µm^2^ field of view. The power density is 60 kW/cm^2^ for single protein imaging. A flow cell for sample delivery is constructed as previously reported^17,18^.

### Protein binding analysis

To track the protein binding and extract each protein binding intensity, we use the Trackmate^23^ plugin in the ImageJ app and a homemade Matlab code with the following settings: 1) In Trackmate, a Laplacian of Gaussian Filter with 8 pixels diameter is set as determined by setup resolution and camera pixel size. 2) A threshold provides a spot with a larger than three signal-to-noise-ratio is set. Here we use 1.5 as the quality threshold in Trackmate. 3) As a result of moving-averaging-differential processing, a binding event will show as a still bright spot with a first-increase-then-decrease intensity profile in a temporal differential image sequence (Figure 1f). Trackmate is used to track the binding intensity temporal changes. A gap tolerance is set to 1 frame, and linking distance is set to 1 pixel for tracking one protein binding. We use Matlab to do the following steps. 4) Extract track information from the Trackmate. Filter out tracks with less than 300 ms residence time. This step effectively removes tracks from the background noise. 5) A frame containing more than 40 spots is also filtered out, which usually originated from one or more large impurity particles miss identified by Trackmate as multiple binding spots. 6) For each track, we use the maximum intensity and the Trackmate quality for protein identification. 7) Separate quality thresholds are determined for different protein species relying on binding quality from Trackmate. We have threshold ranges for anti-IgA (1.5-2.8), IgA (2.8-3.2), and IgM (4.8-10), which are determined in the calibration experiments. The threshold range for the complex is 3.2-4.8.

### IgA and anti-IgA binding kinetics analysis

We have anti-IgA and IgA landing counts as the function of time. The fraction of monomer IgA counts equal to the mole fraction of the solution monomer:

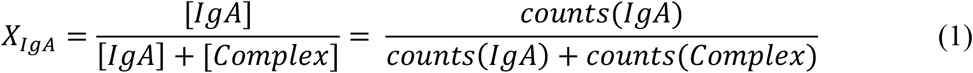

Where [IgA] and [Complex] are the concentration of the IgA and the complex. The sum of IgA and complex concentration is a constant, which is the initial concentration of IgA. We consider the first-order reaction of anti-IgA and IgA association experiment^11,24,25^. According to the derivative equation, *k*_on_ and *k*_off_ can be obtained by fitting the IgA concentration decay:

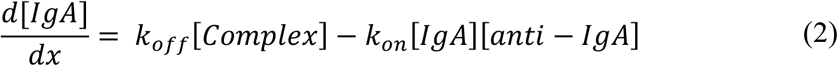

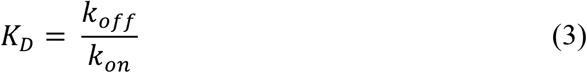

Where *k*_on_ and *k*_off_ are the rate constants, and *K*_D_ is the equilibrium dissociation constant. IgA and complex counts are used here because they have a better signal-to-noise ratio than anti-IgA due to their larger size.

The code is available at https://doi.org/10.5281/zenodo.6583374.

### Surface Functionalization and Surface Immobilization

To measure the nonspecific landing of single proteins, ITO chips were modified with carboxyl groups by the flowing steps. 1) Incubated the ITO chip in NH_3_, H_2_O_2_, and H_2_O mixture with a 1:1:5 volume ratio for 1 h. Dropping water became a layer on the hydrophilic chip surface. 2) Ultrasonically cleaned the chip and container with water, rinsed the chip with water, and dried it with nitrogen gas. 3) Incubated the chip in 1% APTES in IPA for 2 hours. This step functionalizes the surface with the amine group. 4) Ultrasonically cleaned the chip and container with IPA, rinsed the chip with water, and dried it with nitrogen gas. 5) Incubated the chip in 10 g/L succinic anhydrate for 3 hours. The solution pH should be between 7.5-8. 6) Cleaned the chip and container with water ultrasonically, rinsed the chip with water, and dried it with nitrogen gas. Store the chips in the water for further usage. In the experiment, 0.05 M NHS and 0.2 M EDC were mixed and incubated on the chip surface for 15 minutes to activate the surface. The EDC/NHS solution was filtered with a 0.22 μm filter before usage. After activation, the surface was rinsed with PBS. In the calibration experiment, 10 nM IgG or 5 nM IgA in PBS was flowed onto the surface for single protein calibration. In IgM calibration measurement, 20 nM anti-IgM was incubated on the surface for 1 h after EDC/NHS activation. Then, the surface was rinsed with PBS. Finally, 20 nM IgM solution was flowed onto the surface for measurement. In free solution IgA and anti-IgA binding kinetics measurement, 100 nM BSA was incubated on the surface for 1h after EDC/NHS activation. 2.5 nM anti-IgA and 2.5 nM IgA was flowed over the surface. In IgM and anti-IgM binding kinetics experiment, 20 nM anti-IgM was incubated on the surface for 1h after EDC/NHS activation. 2.5 nM IgM solution was flowed onto the surface for measurement. In parallel binding kinetics measurement, 20 nM anti-IgM was incubated on the surface for 1h after EDC/NHS activation. Anti-IgA, IgA, and IgM with a final 2.5 nM concentration for each protein were flowed onto the surface.

## References

1. Moffat, J. G., Vincent, F., Lee, J. A., Eder, J. & Prunotto, M. Opportunities and challenges in phenotypic drug discovery: an industry perspective. Nat. Rev. Drug. Discov. 16, 531–543 (2017).

2. Sadelain, M., Rivière, I. & Riddell, S. Therapeutic T cell engineering. Nature 545, 423–431 (2017).

3. Im, H. et al. Label-free detection and molecular profiling of exosomes with a nano-plasmonic sensor. Nat. Biotechnol. 32, 490–495 (2014).

4. Aćimović, S. S. et al. LSPR Chip for Parallel, Rapid, and Sensitive Detection of Cancer Markers in Serum. Nano Lett. 14, 2636–2641 (2014).

5. Wee, P. & Wang, Z. Epidermal Growth Factor Receptor Cell Proliferation Signaling Pathways. Cancers 9, 52 (2017).

6. Dibble, C. C. & Cantley, L. C. Regulation of mTORC1 by PI3K signaling. Trends in Cell Biology 25, 545–555 (2015).

7. Gingras, A.-C., Gstaiger, M., Raught, B. & Aebersold, R. Analysis of protein complexes using mass spectrometry. Nat. Rev. Mol. Cell. Biol. 8, 645–654 (2007).

8. Jain, A. et al. Probing cellular protein complexes using single-molecule pull-down. Nature 473, 484–488 (2011).

9. Barrios-Rodiles, M. et al. High-Throughput Mapping of a Dynamic Signaling Network in Mammalian Cells. Science 307, 1621–1625 (2005).

10. Campbell, C. & Kim, G. SPR microscopy and its applications to high-throughput analyses of biomolecular binding events and their kinetics. Biomaterials 28, 2380–2392 (2007).

11. Schasfoort, R. B. Handbook of surface plasmon resonance. (Royal Society of Chemistry, 2017).

12. Young, G. et al. Quantitative mass imaging of single biological macromolecules. Science 360, 423–427 (2018).

13. Piliarik, M. & Sandoghdar, V. Direct optical sensing of single unlabelled proteins and super-resolution imaging of their binding sites. Nat. Commun. 5, 4495 (2014).

14. Soltermann, F. et al. Quantifying Protein–Protein Interactions by Molecular Counting with Mass Photometry. Angew. Chem. 132, 10866–10871 (2020).

15. Boyer, D., Tamarat, P., Maali, A., Lounis, B. & Orrit, M. Photothermal Imaging of Nanometer-Sized Metal Particles Among Scatterers. Science 297, 1160–1163 (2002).

16. Selmke, M., Braun, M. & Cichos, F. Photothermal Single-Particle Microscopy: Detection of a Nanolens. ACS Nano 6, 2741–2749 (2012).

17. Zhang, P. et al. Plasmonic scattering imaging of single proteins and binding kinetics. Nat. Methods 17, 1010–1017 (2020).

18. Zhang, P., Ma, G., Wan, Z. & Wang, S. Quantification of Single-Molecule Protein Binding Kinetics in Complex Media with Prism-Coupled Plasmonic Scattering Imaging. ACS Sens. 6, 1357–1366 (2021).

19. Zhang, P. et al. Label-Free Imaging of Nanoscale Displacements and Free-Energy Profiles of Focal Adhesions with Plasmonic Scattering Microscopy. ACS Sens. 6, 4244–4254 (2021).

20. Zhang, P. & Wang, S. Real-Time analysis of exosome secretion of single cells with single molecule imaging. BIOCELL 45, 1449–1451 (2021).

21. Zhang, P. et al. Evanescent scattering imaging of single protein binding kinetics and DNA conformation changes. Nat. Commun. 13, 2298 (2022).

22. Cole, D., Young, G., Weigel, A., Sebesta, A. & Kukura, P. Label-Free Single-Molecule Imaging with Numerical-Aperture-Shaped Interferometric Scattering Microscopy. ACS Photonics 4, 211–216 (2017).

23. Tinevez, J.-Y. et al. TrackMate: An open and extensible platform for single-particle tracking. Methods 115, 80–90 (2017).

24. Bernetti, M., Masetti, M., Rocchia, W. & Cavalli, A. Kinetics of Drug Binding and Residence Time. Annu. Rev. Phys. Chem. 70, 143–171 (2019).

25. Vauquelin, G. Effects of target binding kinetics on in vivo drug efficacy: k_off_, k_on_ and rebinding: Exploring drug rebinding in vivo. British Journal of Pharmacology 173, 2319–2334 (2016).

