## Supplementary material for "Multiplexed protein detection and parallel binding kinetics analysis with label-free digital single-molecule counting": Figure S

**This PDF file includes:**

Figs. S1 to S4
References

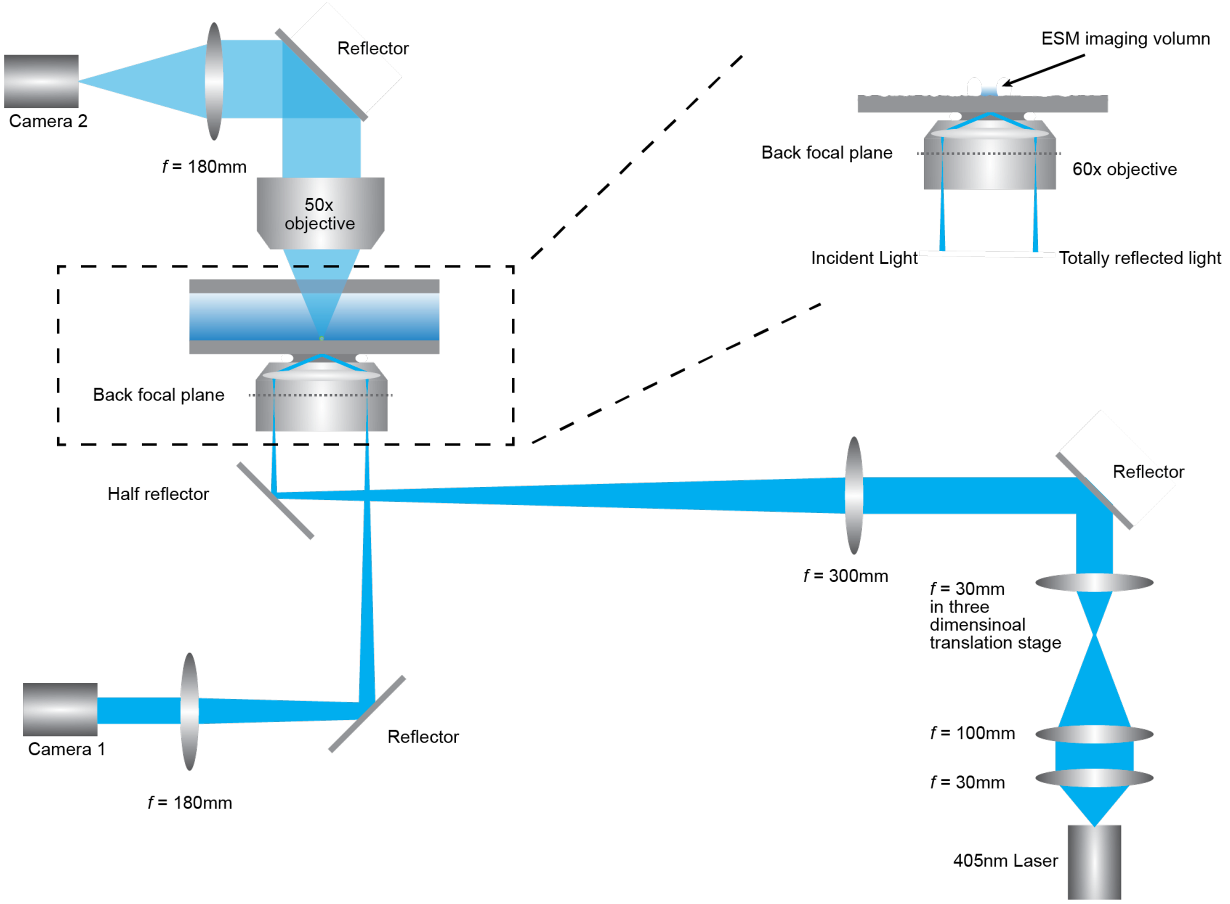


**Figure S1. ESM imaging setup.**

The configuration of ESM is similar to our previous report[1]. 405 nm laser diode (PL450B, Thorlabs, Newton, NJ, US) provides light source through the system. Light beam size is adjusted by a 30mm and a100mm tube lens and incident to the chip at ~65° using a three-dimensional translation stage (XR25P-K2, Thorlabs). We use a top-mounted 50x objective (NA = 0.42) collect scattering light. The camera 2 (MQ003MG-CM, XIMEA) records the scattering signal. Camera 1(MQ013MG-ON, XIMEA) detects the reflection signal.

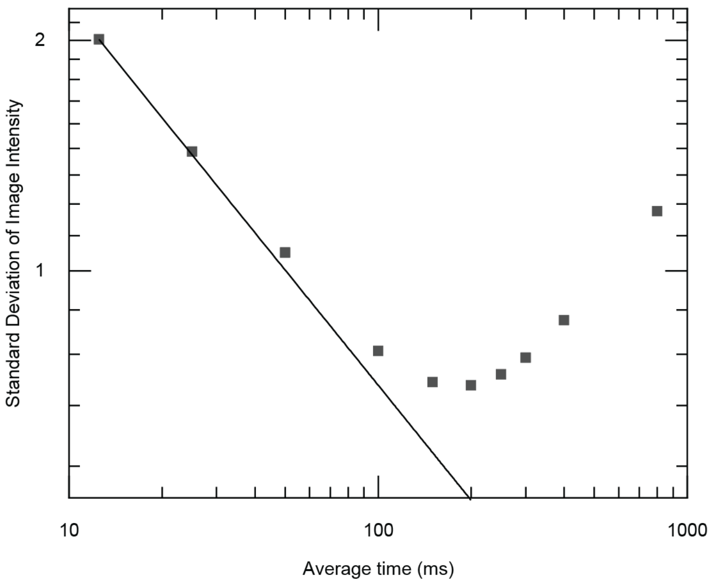


**Figure S2. Standard deviation of processed images as a function of average time.**

The raw images are recorded at 160 fps. Total intensity values in a selected region of interest (~8 pixels in diameter) are obtained at different average time. The dots represent experimental noise at different average time. The solid line is shot noise determined.

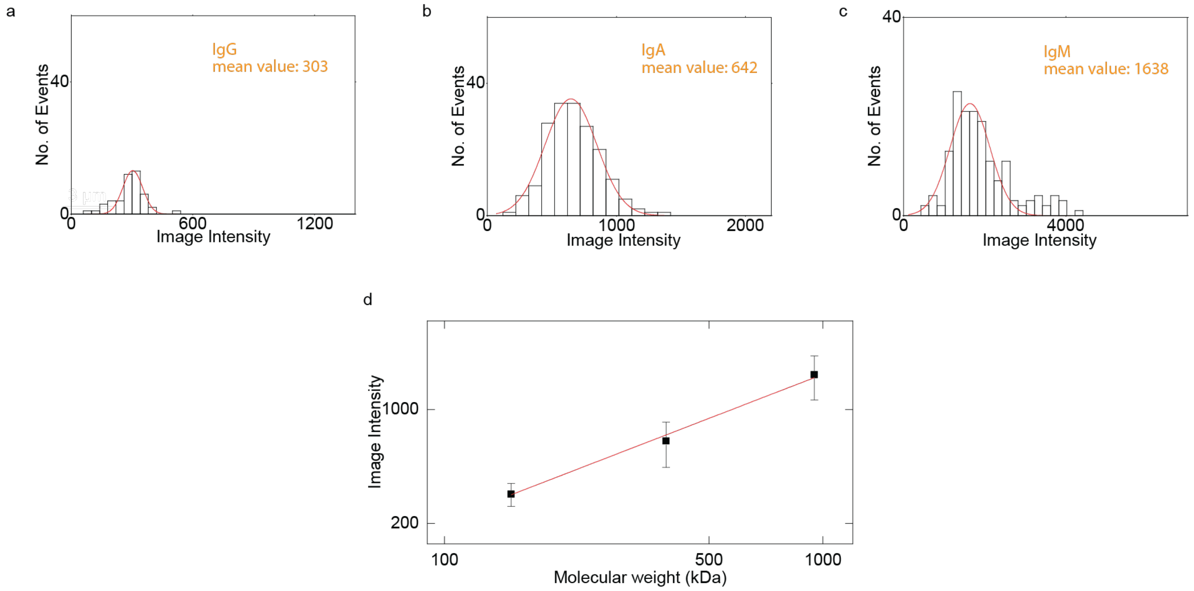


**Figure S3. Calibration result with non-moving average data processing.**

(a-c) IgG, IgA, and IgM images (after image processing) and intensity histogram. The data source is the same as in Figure 2. (d) ESM intensity versus protein mass. The image intensities are obtained from the histograms of each protein in (a-c). The error bars indicate the standard deviation of Gaussian fitting. The red sold line is the linear fitting result.

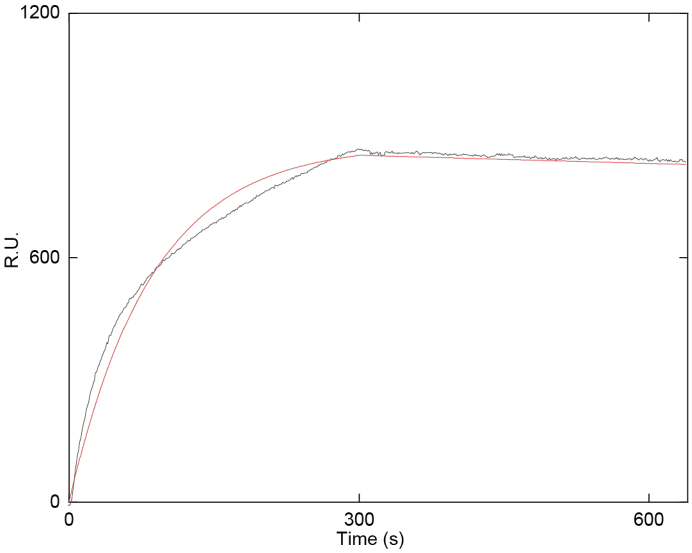


**Figure S4. SPR measurement of IgM binding to anti-IgM.**

50 nM IgM solution was injected into flow chamber with anti-IgM modified on the goat covered cover glass. Black line is experimental result with association and dissociation period. The red line is the fitting result. We obtained $K_D$ = 221 pM, with $k_{on} = 2.77 \times 10^5\ M^{-1}\ s^{-1}$ and $k_{off} = 6.13 \times 10^{-5}\ s^{-1}$. The measurement was performed on a SPR prism setup. The detailed description of the prism SPR will be published in an upcoming article.

### References

1. Zhang, P. *et al.* Evanescent scattering imaging of single protein binding kinetics and DNA conformation changes. *Nat. Commun.* **13**, 2298 (2022).
